# Bioinformatic correction of bacterial morphology-based extraction bias and chimeras in microbiome sequencing data

**DOI:** 10.1101/2023.07.06.547990

**Authors:** Luise Rauer, Amedeo De Tomassi, Christian L. Müller, Claudia Hülpüsch, Claudia Traidl-Hoffmann, Matthias Reiger, Avidan U. Neumann

## Abstract

**Introduction:** Microbiome amplicon sequencing data are distorted by multiple protocol-dependent biases, originating from bacterial DNA extraction, contamination, sequence errors, and chimeras. In particular, extraction bias is a major confounder in sequencing-based microbiome analyses, with no correction method available to date. Here, we suggest using mock community controls to bioinformatically correct extraction bias based on morphological properties.

**Methods:** We compared dilution series of 3 mock communities with an even or staggered composition. DNA was extracted with 8 different extraction protocols (2 buffers, 2 extraction kits, 2 lysis conditions). Extracted DNA was sequenced (V1-V3 16S rRNA gene) together with corresponding DNA mocks. Sequences were denoised using DADA2, and annotated by matching against mock reference genomes.

**Results:** Microbiome composition was significantly different between extraction kits and lysis conditions, but not between buffers. Independent of the extraction protocol, chimera formation increased with high input cell number. Contaminants originated mostly from buffers, and considerable cross-contamination was observed in low-input samples. Comparison of microbiome composition of the cell mocks to corresponding DNA mocks revealed taxon-specific protocol-dependent extraction bias. Strikingly, this extraction bias per species was predictable by bacterial cell morphology. Morphology-based bioinformatic correction of extraction bias significantly improved sample compositions when applied to different samples, even with different taxa.

**Conclusions:** Our results indicate that higher DNA density increases chimera formation during PCR amplification. Furthermore, we show that bioinformatic correction of extraction bias is feasible based on bacterial cell morphology.

## Introduction

Investigating the human microbiome with 16S targeted sequencing has become integral to environmental and medical scientific studies. Recently, awareness has increased that microbiome associated findings lack reproducibility. This inconsistency in findings can be attributed to the wide variety in methods available for generating sequencing-based microbiome data. Each experimental step along the data generation pipeline from sampling to sequencing slightly distorts the data, multiplying into significant protocol-dependent biases [1] that limit the comparability between microbiome studies [2].

One of the major confounders among these biases is extraction bias [3–5], covering the process of cell lysis and isolation of bacterial DNA. Numerous studies have compared extraction methods and found striking differences between extraction kits, preservative buffers, cell lysis approaches, and protocol details like bead size [6] or bead-beating intensity [7]. Traditionally, these differences in extraction methods have been evaluated in terms of cell lysis efficiency, DNA yield, DNA purity, DNA integrity, reproducibility of results, and species richness [8, 9], mostly using environmental samples [9]. However, the key aspect of a protocol’s ability to accurately reflect the original sample composition can only be assessed with the help of standardized controls like mock communities. Such standardized mock communities are increasingly developed [10–12] and used, e.g., to evaluate extraction bias [5, 13, 14].

Mock communities also allow for evaluating sthe impact of other spurious sequences, such as contaminants, amplification and sequencing errors, and chimeras. Contaminants frequently originate from lab reagents and operators during DNA extraction and PCR amplification [15]. Besides external DNA contamination, internal cross-contamination by index hopping and bleed-through may significantly blur microbial signatures but is rarely investigated [16]. Despite laboratory efforts, both contamination and cross-contamination remain particularly problematic for low biomass samples [15–18], such as from milk, lung, or skin microbiomes [19].

Similarly, amplification and sequencing artifacts such as sequence errors and chimeras can critically impact taxonomic annotation and inflate diversity estimates. While modern high-fidelity proof-reading polymerases report very low error rates per sequence doubling, sequence errors may still accumulate over multiple PCR cycles and require clustering or denoising of sequences before biological interpretation [20]. Chimera formation remains an inherent problem in multi-template PCR reactions with high homology between templates, as in 16S sequencing experiments.

While computational pipelines can be used to correct sequence errors (e.g., DADA2 [20], deblur [21]), remove chimeras (e.g., UCHIME [22], ChimeraSlayer [23]), or remove contaminants (e.g., Decontam [24], MicrobIEM [25]), there is currently no laboratory or bioinformatic approach to overcome extraction bias. Moreover, these algorithms mainly tackle individual biases, whereas mock communities allow for analyzing the origin and interaction of bias accumulated over several protocol steps [1]. We designed this study to investigate extraction bias and its relation to the morphological properties of bacterial cells. In order to accurately quantify extraction bias, we also explore downstream protocol biases from (cross-)contamination, sequence errors, and chimera formation. Our analyses indicate that mock communities may be used to measure and bioinformatically correct differential DNA extraction efficiencies and other biases in microbiome sequencing data, paving the road to cross-protocol comparisons in microbiome research.

## Methods

### Study design and sample preparation

#### Mock community samples and negative controls

An overview of our study design is shown in **Figure 1**. We used mock microbial community standards with different taxa and abundance compositions provided by the ZymoBIOMICS series of ZymoResearch (Freiburg, Germany). Two of these mock communities contain the eight bacterial species *Bacillus subtilis*, *Lactobacillus fermentum* (also known as *Limosilactobacillus fermentum*), *Pseudomonas aeruginosa*, *Salmonella enterica*, *Escherichia coli*, *Listeria monocytogenes*, and *Enterococcus faecalis*, and the two fungal species *Saccharomyces cerevisiae* and *Cryptococcus neoformans*. The latter two cannot be detected by our 16S rRNA gene sequencing approach and are therefore not further discussed. This eight-bacteria mock is available as a whole-cell mock community with an even (D6300) or staggered composition (D6310), and as corresponding DNA mocks with an even (D6305) or staggered (D6311) composition of DNA. The spike-in community (D6321) is only available as a staggered whole-cell community and consists of the three species *Imtechella halotolerans*, *Allobacillus halotolerans*, and *Truepera radiovictrix*, all of which are alien to the human microbiome. **Table 1** shows the expected composition of all mock communities, considering 16S copy numbers and genome size for 16S rRNA gene sequencing experiments.

**Figure 1.**
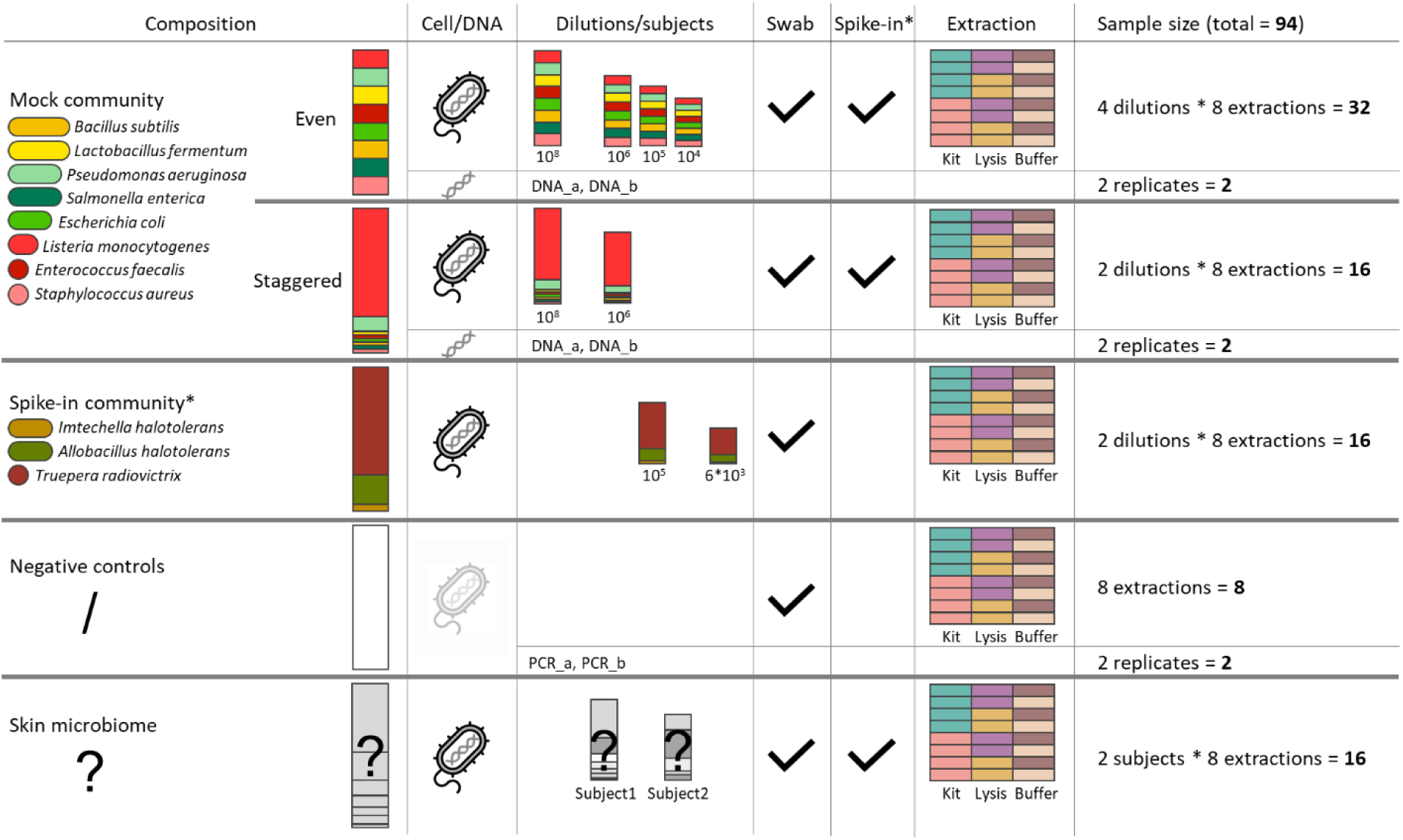
Study design of mock communities, controls and skin microbiome samples to study extraction bias in eight extraction protocols. Cell-based mock communities with an even or a staggered composition diluted to 10^8^ to 10^4^ input cells and their corresponding DNA mocks were used to determine extraction bias. Eight different extraction protocols were tested in a combination of two extraction kits, two lysis protocols, and two extraction buffers. Additionally, negative controls and environmental skin microbiome swab samples underwent these eight protocols. A swab was also added to all non-skin samples. Asterisks (*) denote the spike-in community, of which 6*10^3^ cells were spiked into each mock and skin sample, and of which pure spike-in community samples were also investigated in two dilutions from 10^5^ to 6*10^3^ input cells.

**Table 1:**
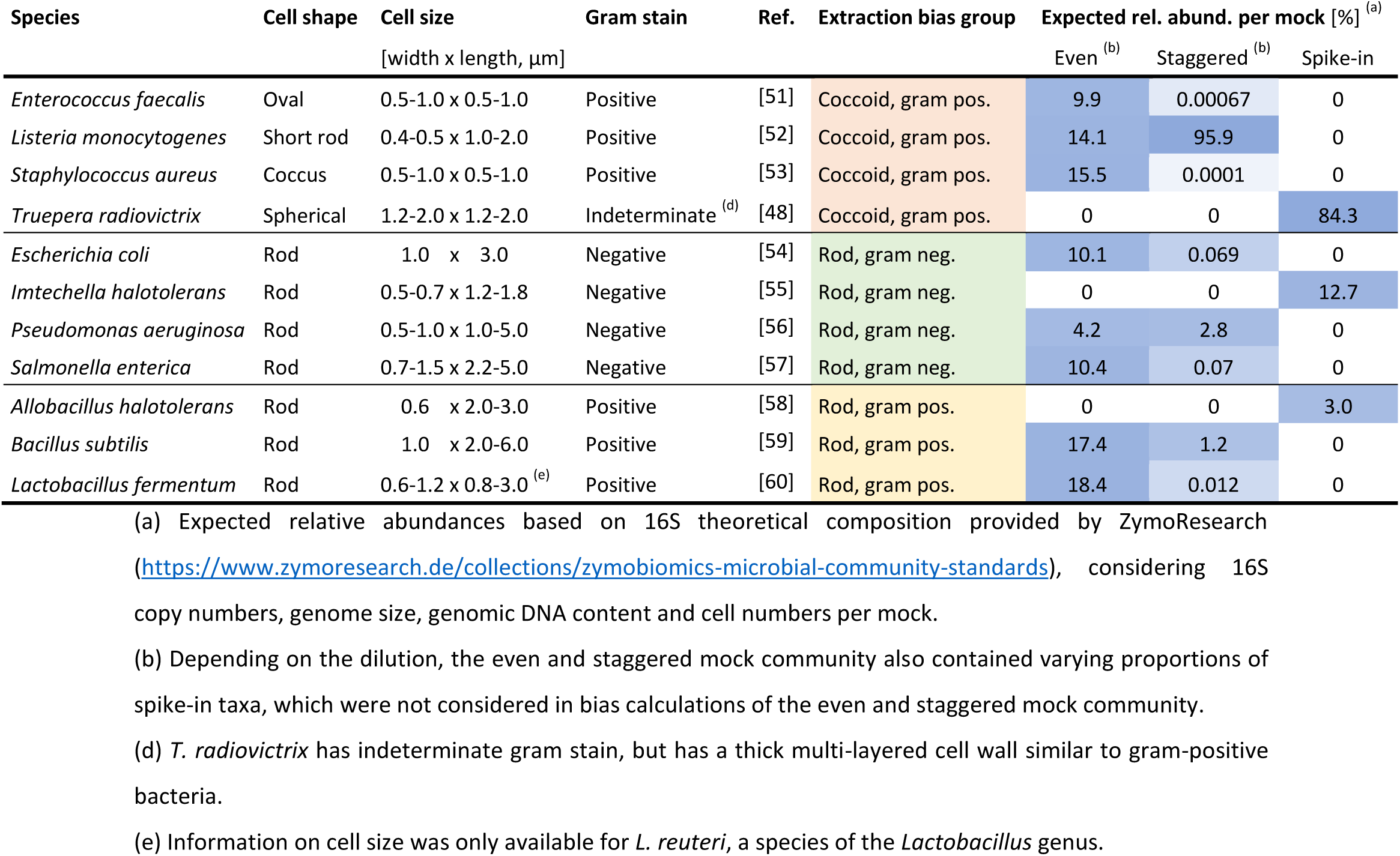
Based on cell shape and gram stain, bacterial taxa can be classified into three morphological groups that determine extraction bias in mock data. Cell shape, cell size, and gram stain were extracted from scientific references (Ref.), and were used to divide all eleven mock taxa into three morphological groups. Red, green, and yellow background color highlight extraction bias groups, darker blue background color indicates higher expected relative abundance (rel. abund.) per mock community.

| Species | Cell shape | Cell size<br>[width x length, µm] | Gram stain | Ref. | Extraction bias group | Expected rel. abund. per mock [%] <sup>(a)</sup> |  |  |
| --- | --- | --- | --- | --- | --- | --- | --- | --- |
|  |  |  |  |  |  | Even <sup>(b)</sup> | Staggered <sup>(b)</sup> | Spike-in |
| *Enterococcus faecalis* | Oval | 0.5-1.0 x 0.5-1.0 | Positive | [51] | Coccoid, gram pos. | 9.9 | 0.00067 | 0 |
| *Listeria monocytogenes* | Short rod | 0.4-0.5 x 1.0-2.0 | Positive | [52] | Coccoid, gram pos. | 14.1 | 95.9 | 0 |
| *Staphylococcus aureus* | Coccus | 0.5-1.0 x 0.5-1.0 | Positive | [53] | Coccoid, gram pos. | 15.5 | 0.0001 | 0 |
| *Trueperia radiovictrix* | Spherical | 1.2-2.0 x 1.2-2.0 | Indeterminate <sup>(d)</sup> | [48] | Coccoid, gram pos. | 0 | 0 | 84.3 |
| *Escherichia coli* | Rod | 1.0 x 3.0 | Negative | [54] | Rod, gram neg. | 10.1 | 0.069 | 0 |
| *Imtechella halotolerans* | Rod | 0.5-0.7 x 1.2-1.8 | Negative | [55] | Rod, gram neg. | 0 | 0 | 12.7 |
| *Pseudomonas aeruginosa* | Rod | 0.5-1.0 x 1.0-5.0 | Negative | [56] | Rod, gram neg. | 4.2 | 2.8 | 0 |
| *Salmonella enterica* | Rod | 0.7-1.5 x 2.2-5.0 | Negative | [57] | Rod, gram neg. | 10.4 | 0.07 | 0 |
| *Allobacillus halotolerans* | Rod | 0.6 x 2.0-3.0 | Positive | [58] | Rod, gram pos. | 0 | 0 | 3.0 |
| *Bacillus subtilis* | Rod | 1.0 x 2.0-6.0 | Positive | [59] | Rod, gram pos. | 17.4 | 1.2 | 0 |
| *Lactobacillus fermentum* | Rod | 0.6-1.2 x 0.8-3.0 <sup>(e)</sup> | Positive | [60] | Rod, gram pos. | 18.4 | 0.012 | 0 |

(a) Expected relative abundances based on 16S theoretical composition provided by ZymoResearch (https://www.zymoresearch.de/collections/zymobiomics-microbial-community-standards), considering 16S copy numbers, genome size, genomic DNA content and cell numbers per mock.
(b) Depending on the dilution, the even and staggered mock community also contained varying proportions of spike-in taxa, which were not considered in bias calculations of the even and staggered mock community.
(d) *T. radiovictrix* has indeterminate gram stain, but has a thick multi-layered cell wall similar to gram-positive bacteria.
(e) Information on cell size was only available for *L. reuteri*, a species of the *Lactobacillus* genus.

The three cell mock communities (even, staggered, and spike-in) were diluted with Buffer AVE (Qiagen, Hilden, Germany) and split into eight replicates per mock and dilution for subsequent DNA extraction. The final mock bacterial input per sample ranged from 10^8^ to 5.55*10^3^ cells (rounded to 6*10^3^ cells in all figures), as shown in **Figure 1**. All 48 even and staggered cell mock samples were spiked with 5.55*10^3^ cells of the spike-in mock. The 64 cell mock samples (even, staggered, and spike-in) also contained empty swabs (DRYSWAB MWE, Cat No. MW 940/125, Medical Wire, Corsham/Wiltshire, UK). Eight empty tubes containing only a swab were processed along with the samples as full-pipeline negative controls.

#### Skin microbiome samples

The skin microbiome of two healthy subjects was sampled in replicates by striking the forearm with eight parallel swabs (DRYSWAB MWE Cat No. MW 940/125, Medical Wire). These 16 environmental skin microbiome samples were also spiked with 5.55*10^3^ cells of the spike-in mock. Study subjects provided written informed consent of participation.

### Sample processing

#### DNA extraction

The eight replicates of each mock dilution and subject underwent eight different extraction protocols, representing a combination of two extraction kits, two lysis conditions, and two extraction buffers. We compared the extraction kits QIAamp UCP Pathogen Mini Kit (Cat No. 50214, Qiagen, Hilden, Germany; ‘Q’) versus ZymoBIOMICS DNA Microprep Kit (Cat No. D4301, ZymoResearch, Freiburg, Germany, ‘Z’). The kit variable includes the beads used for cell lysis, with 0.1- and 0.5-mm Zirconia beads provided in the ZymoResearch kit (‘Z’) and 0.1 µm Zirconia Beads (BioSpec, Bartlesville, Oklahoma) used for the Qiagen kit (‘Q’). The two lysis conditions were a ‘soft’ lysis protocol (‘S’) at 5.6 KRPM for 3 min versus a rather ‘tough’ lysis protocol (‘T’) at 9.0 KRPM for 4 min, both on a Precellys Evolution Touch homogenizer (Bertin, Montigny-le-Bretonneux, France). Extraction buffers comprised DNA/RNA shield R1100-50 stabilizer (‘z’) provided with the ZymoResearch kit, and a combination of Stool Stabilizer (Stratec, Birkenfeld, Germany) as preservative and Buffer ATL provided in the Qiagen kit (both summarized as ‘q’). Abbreviations per extraction protocol combination are used throughout this work in the format ‘kit_lysis_buffer’, e.g., ‘Q_T_z’ for the combination of the Qiagen extraction kit, the tough lysis protocol, and the ZymoResearch buffer.

All 88 whole-cell samples of diluted mock, full-pipeline negative controls, and skin microbiome with their respective buffers were frozen before further processing, and only the Buffer ATL of ‘q’ was added after thawing the samples for ensuing DNA extraction.

#### 16S rRNA gene amplification and sequencing

After extraction, we added two replicates of even (D6305) and staggered (D6311) DNA mock community samples with 0.1µl/ng DNA concentration (roughly equivalent to 1.3*10^7^ input cells), and two PCR negative controls, summing to a total of 94 samples. In a first PCR step, the V1-V3 variable region of the 16S rRNA gene was amplified using the Q5 Hot Start High-Fidelity DNA polymerase (New England Biolabs, Ipswich, Massachusetts, USA), and primers 27F-YM (5’-AGAGTTTGATYMTGGCTCAG-3’) and 534R (5’-ATTACCGCGGCTGCTGG-3’) with Illumina adaptor sequences. PCR conditions included an initial denaturation step at 98°C for 1 min, followed by 25 cycles of 98°C for 10s, 59°C for 20s, and 72°C for 15s, and a final elongation step at 72°C for 2 min. In a second PCR step, sample-specific dual indexed barcodes were added to allow for multiplexed sequencing. The second PCR was run with an initial denaturation step at 98°C for 40s, followed by 8 cycles of 98°C for 20s, 55°C for 40s, and 72°C for 40s, and a final elongation step at 72°C for 2 min. Indexed amplicons were purified twice using AMPure XP beads (Beckman Coulter, Fullerton, California, USA) according to manufacturer’s instructions and quantified using the fluorescent dye-based Qubit® dsDNA HS Assay Kit (Invitrogen, Carlsbad, California, USA). Samples were equimolarly pooled and sequenced using the Illumina MiSeq platform (Illumina Inc., San Diego, California, USA) with the MiSeq® Reagent Kit v3 (Illumina Inc.) to produce 2×300 bp reads.

### Data analysis

#### Raw sequence processing

High-quality reads according to Illumina specifications were de-multiplexed by their sample-specific index barcode sequences using the MiSeq® Reporter software. Raw sequences from FASTQ files were quality-controlled and denoised using DADA2 (version 1.16.0) [20]. Default filtering parameters were used, except for trimLeft = 20 to remove the forward primer and truncLen = 299, and nbases = 10^9^ for learning error rates in DADA2. Due to relatively low-quality reverse reads, we processed only the forward reads to keep the highest possible proportion of sequences. We did not remove chimeras or contaminants to allow for investigation of chimera formation and protocol contaminants.

#### Taxonomic classification

Taxonomic annotation of the resulting ASVs was performed by comparison against the reference 16S gene sequences provided by ZymoResearch (available from https://s3.amazonaws.com/zymo-files/BioPool/ZymoBIOMICS.STD.refseq.v2.zip). Levenshtein distance (edit distance, LV) was calculated between expected sequences (cut to 279 bp) and observed ASV sequences, with LV = 0 indicating an exact match with the reference sequence. We used the smallest LV distance to any expected sequence as a proxy for the number of sequence errors (substitutions or indels) introduced during amplification or sequencing. Ambiguous LV annotations between *E. coli* and *S. enterica* were resolved for 6 ASV sequences with LV ≤ 8 (maximum relative abundance 2.3%) using the DNA evolution model [26]. ASV sequences with LV > 8 to any expected sequence (corresponding to < 97 % identity) were further checked for their longest common substring (LCS) with the 279bp-long expected reference sequences. When > 95 % (266 bp) of the sequence exactly matched with parts of two or three different mock taxa, we interpreted these sequences to represent chimeras between mock taxa. All remaining ASVs that could not be explained by low LV distance or a mixture of exact matches were defined as “Unclassified”, representing mostly skin and contaminant taxa, but potentially also chimeras with ≥ 4 parts or chimeras between mock and skin/contaminant taxa.

Additionally, we used the RDP- and NCBI-based annotation provided through DADA2 [27] to get taxonomic information on species not part of the ZymoResearch mock communities, such as the skin taxa.

#### Statistical analysis

All statistical analyses and visualizations were created using R (version 4.0.2) [28]. The R packages ‘tidyverse’ (version 2.0.0) [29], ‘reshape2’ (version 1.4.4) [30], ‘ggh4x’ (version 0.2.4) [31], ‘ggConvexHull’ (version 0.1.0) [32], and ‘ComplexHeatmap’ (version 2.4.3) [33] were used for data preparation and visualization. Clustering of heatmaps was performed using Euclidean distance and complete linkage. Clustering of ‘Unclassified’ ASVs was done by kmeans() implemented in R with default parameters and 4 clusters. Levenshtein (LV) distance was calculated using the R package ‘stringdist’ (version 0.9.10) [34], the DNA evolution model using the package ‘ape’ (version 5.7-1) [35] and the longest common substring (LCS) using the package ‘PTXQC’ (version 1.0.16) [36].

ASVs with LV ≤ 4 to any expected sequence were accepted to represent mock taxa and were summarized to species level for subsequent analyses. Similarly, DADA2-based taxonomic annotation was summarized to genus level for all analyses of the skin samples. Spearman correlation was used to measure associations between continuous variables. When calculating distances between samples, we used the traditional Bray-Curtis dissimilarity and the compositional Aitchison distance. For all analyses using Bray-Curtis dissimilarities, reads were transformed to relative abundances by total sum scaling (TSS). For all analyses using Aitchison distance or logarithm, zero counts were replaced by 0.5 and zero relative abundances by 0.00001. For beta diversity analyses of mock and spike-in samples, only mock taxa were considered. Bray-Curtis or Aitchison distances were visualized using principal coordinate analysis (PCoA) provided through wcmdscale() of the r package ‘vegan’ (version 2.6-4) [37], and corresponding p-values were calculated using PERMANOVA.

To calculate bias per taxon and protocol, we applied the metacal approach [1] implemented in the R package metacal (version 0.2.0) [38]. Metacal represents a compositional mathematical model, that assumes microbiome sequencing data to be a multiplicative result of input relative abundance and taxon-specific biases, and allows for calculating and correcting these biases. In the staggered mock community, only the four most abundant taxa (with expected relative abundance ≥ 0.07%, see **Table 1**) were consistently detected across the staggered cell and DNA mock samples. Since Aitchison distance is sensitive to changes in small count values, we only considered these four consistently detected taxa in all bias calculations and corrections. For the same reason, we excluded from all metacal-based analyses one 10^4^ sample of the even mock community, where *P. aeruginosa* was not detected, and one 6*10^3^ sample of the spike-in community, where none of the spike-in taxa were detected. For all analyses using randomness, set.seed(1) was run for reproducibility. Reduction in bias after bioinformatic correction was assessed using Wilcoxon signed-rank test.

## Results

### Sequence classification and sequence errors

We prepared dilutions series of three commercially available mock communities from ZymoResearch. These mocks were used together with negative controls and environmental skin samples to study eight different extraction protocols in a combination of two extraction buffers, two lysis protocols, and two extraction kits. The resulting ASVs from V1-V3 16S rRNA gene sequencing were taxonomically annotated by matching against reference sequences provided by ZymoResearch. This taxonomic matching was done by combining LV distance with LCS, which allowed for classifying ASVs into exact matches, sequence errors, chimeras between mock sequences, and remaining ‘Unclassified’ sequences (**Supp. Figure 1A**).

The highest proportion of correct sequences was observed in the staggered mock, reaching 99.7% (median) of exact matches in samples with 10^6^ input cells. All even mock samples presented with considerably larger proportions of sequence errors (range 6.2% to 31.6%, median 22.3%). Interestingly, most sequence error ASVs were only present in the 10^8^ input samples. In contrast, the majority of sequence error reads in even mock samples was assigned to 8 ASVs of *L. fermentum*, *E. coli*, or *S. enterica* (76.1%), and was consistently detected across all even mock samples (**Supp. Figure 2**). These three species jointly make up only 0.15% of the expected staggered mock composition, explaining the lower proportion of sequence errors observed in the staggered mock.

**Figure 2.**
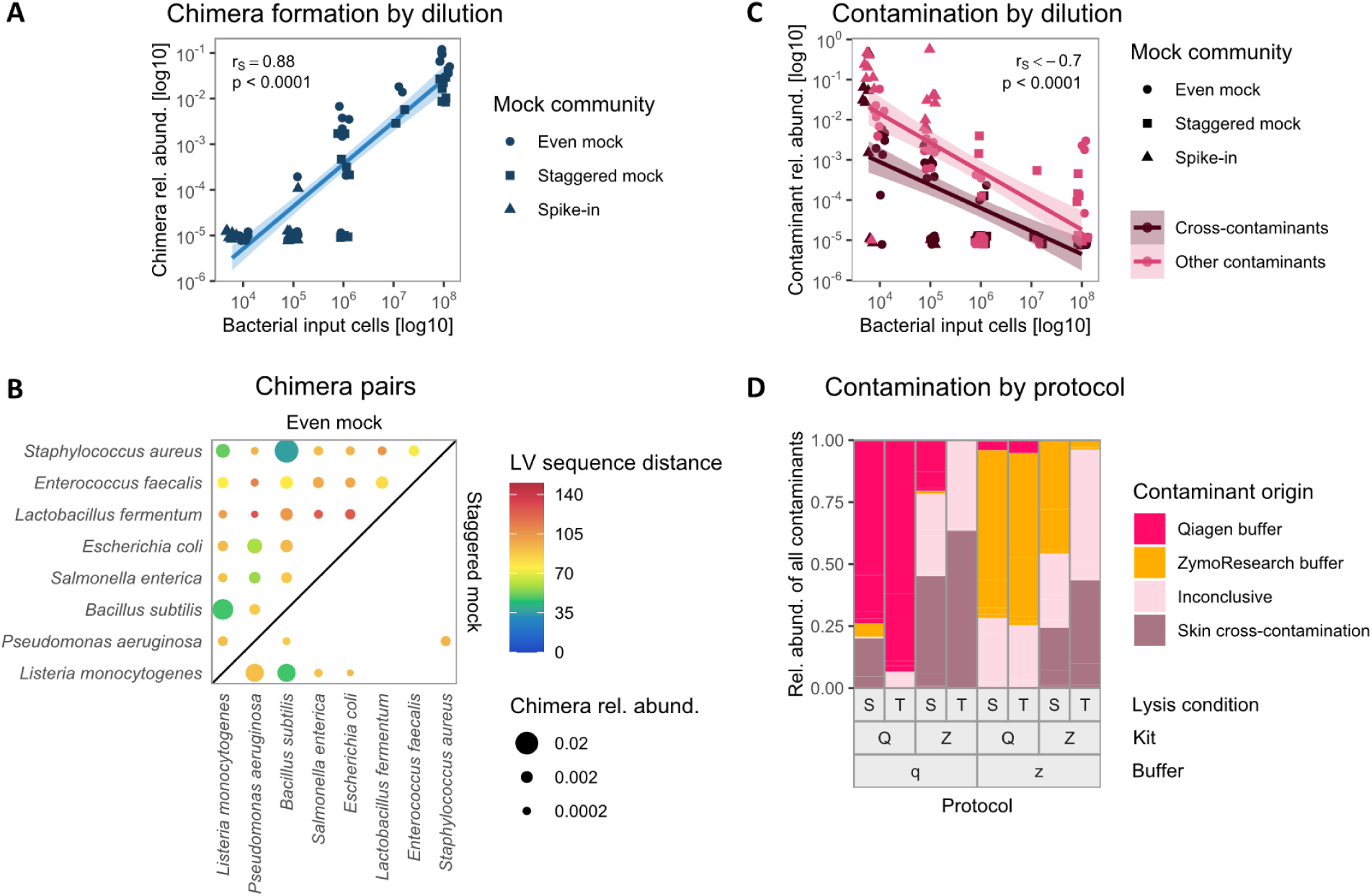
Chimera formation between closely related and highly abundant species increases with high bacterial input, and contamination decreases with bacterial input and originates mostly from buffers. Chimera relative abundance is significantly positively correlated with bacterial input cells per sample across all mock communities (**A**). Chimeras were predominantly formed between closely related species in the even mock (upper left diagonal), indicated by blue/green color, or between highly abundant species in the staggered mock (lower right diagonal), as indicated by species order in decreasing expected abundance from left to right (**B**). Both cross-contaminants and other contaminants significantly negatively correlated with bacterial input cells per sample across all mock communities (**C**). Contamination was substantially associated with extraction protocols, with distinct contaminants originating from the two extraction buffers (**D**). Zeros (**A**, **C**) were replaced by 0.00001. Correlations were estimated by Spearman’s rank correlation coefficient rho (r_S_). Chimera pairs (**B**) are only shown for 10^8^ input cell samples, and point area indicates chimera relative abundance per sample. Chimeras (LV > 8) between *E. coli* and *S. enterica* could not be identified due to their small sequence distance (minimum LV = 6). Contaminant categories (**D**) were determined by kmeans clustering of ASV relative abundances over all mock and control samples, relative abundance per contaminant origin is shown out of total contamination per sample. LV: Levenshtein, Rel. abund.: relative abundance, Q: Qiagen extraction kit, Z: ZymoResearch extraction kit, S: ‘soft’ lysis condition, T: ‘tough’ lysis condition, q: Qiagen/Stratec buffer, z: ZymoResearch buffer.

Due to the ubiquitous presence of sequence errors with LV ≤ 4 in our data (**Supp. Figure 1A**), we accepted these sequences to represent valid mock taxa for all subsequent analyses. The resulting final taxonomic distribution of mock and spike-in taxa over samples is shown in **Supp. Figure 1B**.

### Chimera formation

Surprisingly, chimera formation increased with higher sample biomass, reaching up to 11% in samples with 10^8^ input cells (**Supp. Figure 1A**). Samples with ≤ 10^5^ input cells contained almost no chimeras, and the proportion of chimeric reads was significantly positively correlated with the number of input cells per sample (r_S_ = 0.88, p < 0.001, **Figure 2A**).

Investigating chimera formation in more detail, we found that most chimeric combinations were consistently detected across protocols (**Supp. Figure 3**). Many three-part chimeras (“trimeras”) were actually composed of only two different species, interrupting the exact matching of the LCS with short sequence errors. Therefore, we focused only on the two unique species with the longest LCS per chimera for subsequent analyses, and on samples with 10^8^ input cells due to their high abundance of chimeras. Chimeras in the even mock were overall more diverse, and all discernible taxon combinations were detected (**Figure 2B**). However, most chimeric reads were formed by pairs of *S. aureus*, *B. subtilis*, and *L. monocytogenes*, or of *P. aeruginosa* and *E. coli*., reflecting the groups of taxa with high sequence similarity (**Figure 3B**, **Supp. Figure 4**). In the staggered mock, we observed overall less chimera formation and fewer combinations of taxa, with chimeras mainly formed between the most abundant species *L. monocytogenes*, *B. subtilis*, and *P. aeruginosa* (**Figure 2B**). Taken together, our data indicate that higher microbial DNA input leads to increased chimera formation between closely related and highly abundant sequences.

**Figure 3.**
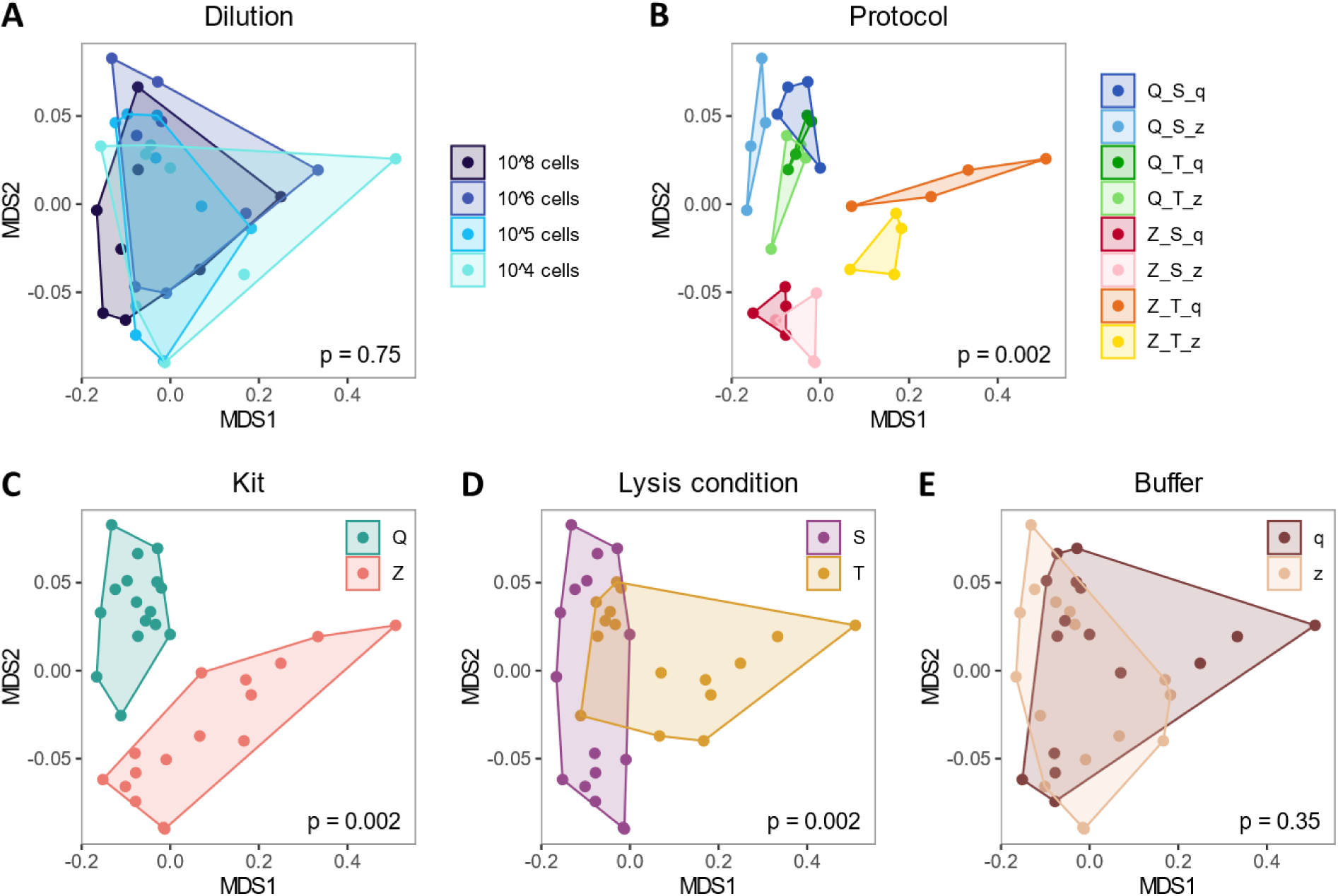
Sample composition is significantly affected by extraction protocols, particularly by extraction kit and lysis condition in the even mock community. Beta diversity analysis revealed significant differences in global mock composition between extraction protocols (**B**), kits (**C**), and lysis conditions (**D**), but not between dilutions (**A**) or buffers (**E**). Beta diversity was performed only on mock taxa with LV ≤ 4 to any expected mock sequence, and is visualized by PCoA on Bray-Curtis dissimilarities. Polygonal shaded areas connect samples of the same group, p-values are derived from PERMANOVA tests with 500 permutations. Q: Qiagen extraction kit, Z: ZymoResearch extraction kit, S: ‘soft’ lysis condition, T: ‘tough’ lysis condition, q: Qiagen/Stratec buffer, z: ZymoResearch buffer.

**Figure 4.**
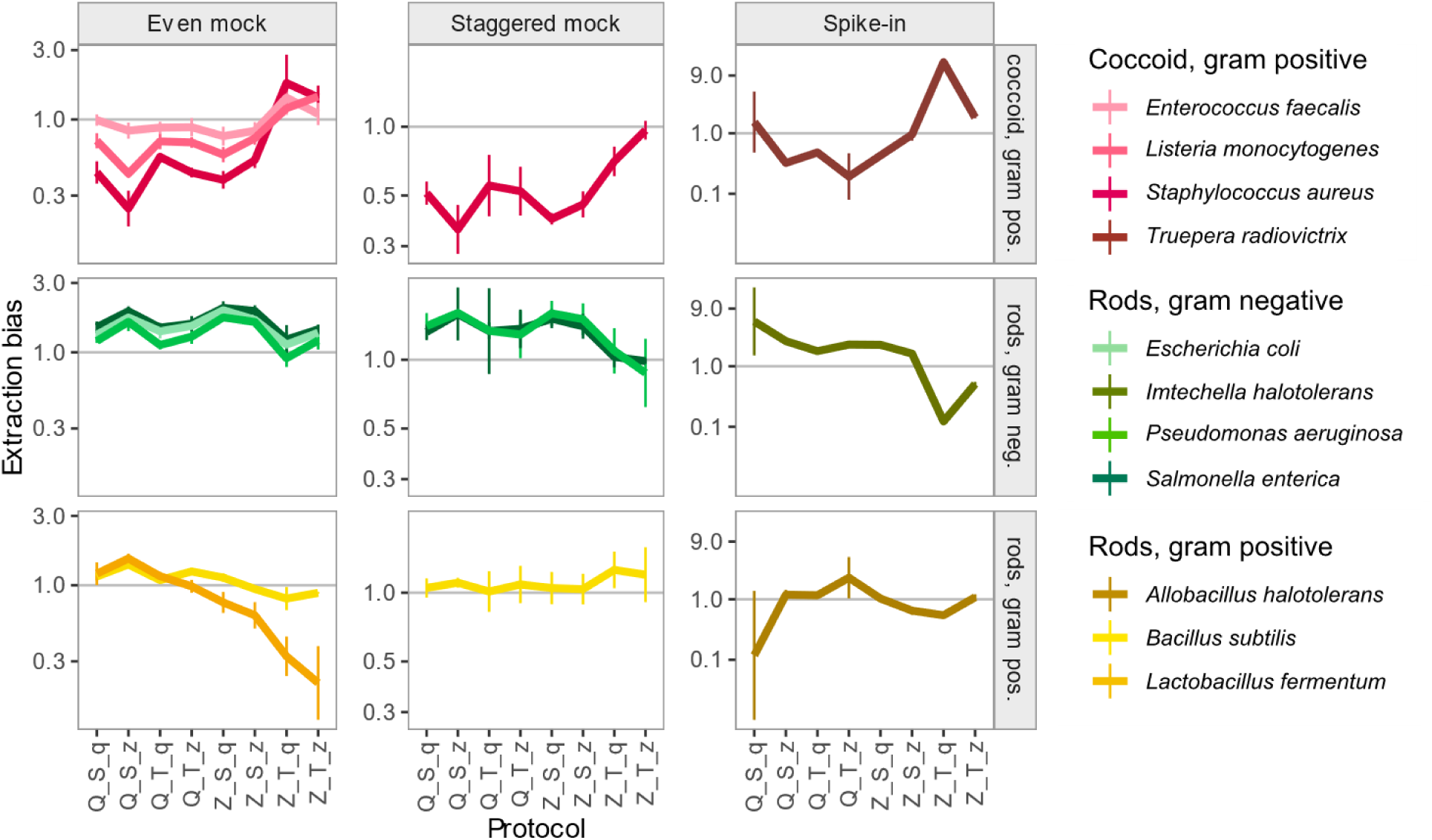
Extraction bias per protocol is highly similar between taxa of the same morphology-based group over all mock communities. Morphological groups of extraction bias were formed based on cell shape and gram stain. Using the metacal R package, extraction bias was calculated per protocol over all dilutions of each mock community in relation the respective DNA mock composition (even, staggered mock) or to the expected mock composition (spike-in mock). Vertical lines represent the standard error per protocol and mock, based on four dilutions (even mock) or two dilutions (staggered mock, spike-in mock). Gram pos.: gram positive, gram neg: gram negative, Q: Qiagen extraction kit, Z: ZymoResearch extraction kit, S: ‘soft’ lysis condition, T: ‘tough’ lysis condition, q: Qiagen/Stratec buffer, z: ZymoResearch buffer.

### Contamination and cross-contamination

As expected, most ASVs in the skin samples did not match the mock reference sequences and were classified as ‘Unclassified’ by LV and LCS matching, apart from small proportions of spike-in taxa spiked into the skin samples. Higher proportions of ‘Unclassified’ ASVs were also found in samples with low bacterial input, such as in the 10^4^ even mock, the 10^5^ and 5.55*10^3^ spike-in dilutions, and negative controls, probably representing contamination or cross-contamination (**Supp. Figure 1A, B**).

We further investigated these presumably contaminating ‘Unclassified’ ASVs present in at least three mock and control samples using kmeans clustering of their relative abundances per sample. Almost half of these ‘Unclassified’ reads (median 48.8%) originated from the two extraction buffers (**Figure 2D**, **Supp. Figure 5**). Using the RDP-based annotation, contaminants present in the Stratec/Qiagen buffer (‘q’) were assigned to *Methylobacterium*, *Alcaligenes*, and *Brucella*, whereas contaminants present in the ZymoResearch buffer (‘z’) belonged to *Paraburkholderia*, *Aquabacterium*, *Nitrospirillum*, and *Herbaspirillum*. Another group of two clustered ASVs was assigned to *Cutibacterium* and *Pseudomonas*. Together with other *Staphylococcus* and *Corynebacterium* ASVs in the fourth cluster of ‘inconclusive’ origin, these four genera were all among the top 10 genera in our skin microbiome samples, indicating contamination from lab operators or cross-contamination from the skin samples. We observed not only skin-to-mock contamination, but also mock-to-skin and mock-to-control contamination (**Supp. Figure 1A, B**), clearly supporting cross-contamination in our data. Overall, the proportion of both contaminants (r_S_ = -0.74, p < 0.0001) and cross-contaminants (r_S_ = -0.70, p < 0.0001) significantly decreased with higher bacterial input cell numbers (**Figure 2C**).

**Figure 5.**
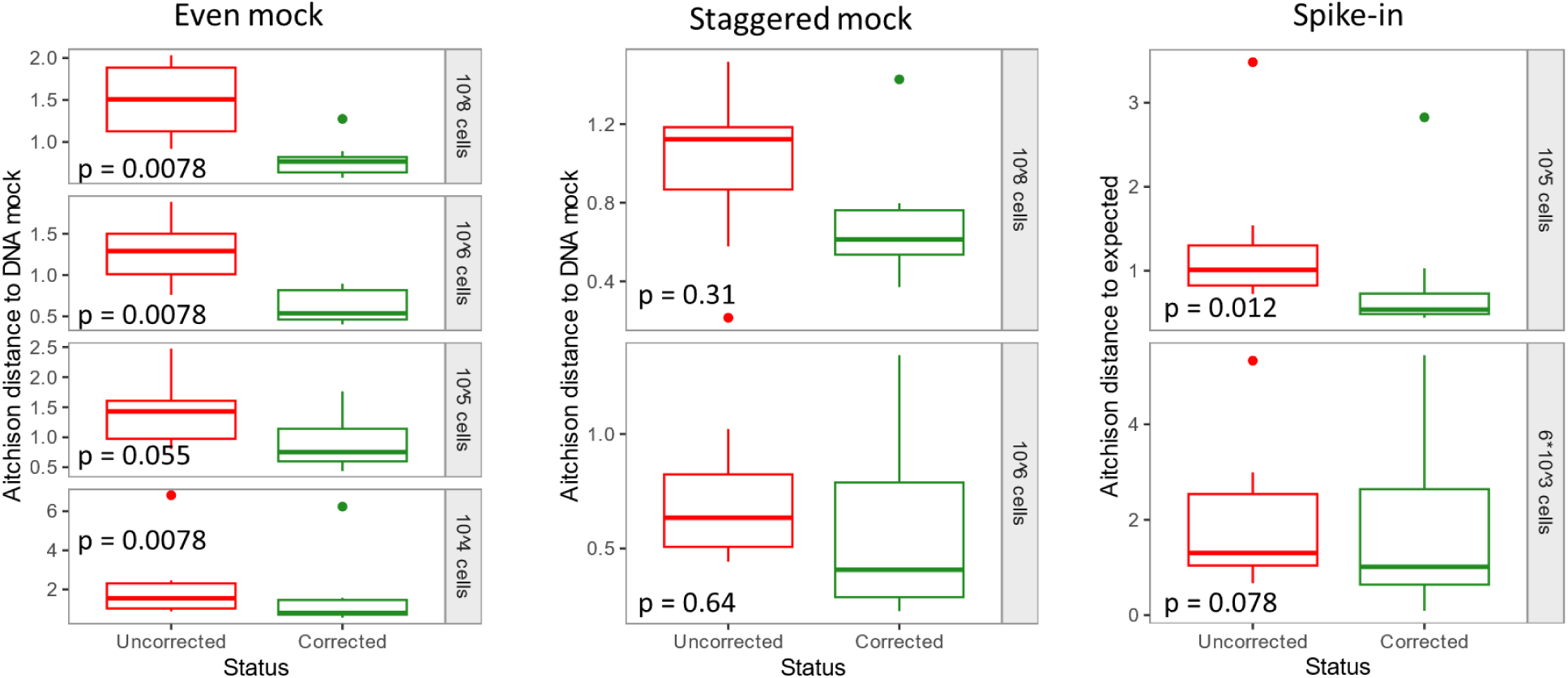
Distance to the DNA or expected mock composition is reduced after applying morphology based correction of extraction bias per extraction protocol in all mock communities. Extraction bias per protocol was calculated from the 10^6^ even mock samples, summarized by bacterial morphology group, and applied to the 10^6^ even mock sample (internal correction), but also to different samples of the same mock (even mock 10^8^, 10^5^, 10^4^ cells), different samples of a different mock (staggered mock), and to different samples with different taxa (spike-in mock). Extraction bias was measured as Aitchison distance to the DNA mock composition (even mock, staggered mock) or as distance to the expected mock composition (spike-in mock). Boxes denote the median and interquartile range (IQR), whiskers represent values up to 1.5 times the IQR, dots indicate samples outside the IQR.

### Effect of extraction protocols

Having resolved the origin of sequence errors, chimeras and contaminants, we investigated the effect of extraction protocols on global sample composition, focusing only on each sample’s expected taxa with LV ≤ 4 in the following beta diversity analyses.

Considering the eight expected bacteria in the even mock, we found no significant difference in sample composition between dilutions (p = 0.75 BC/0.85 Aitchison, **Figure 3A**). Therefore, we treated dilutions as replicates for studying protocol effects (**Figure 3B-E**). The choice of our eight protocols had substantial effects on sample composition, creating eight almost distinct groups of samples (p = 0.0020 BC/Aitchison, **Figure 3B**). When comparing protocol details, we found significant differences between extraction kits (p = 0.0020 BC/Aitchison, **Figure 3C**) and lysis conditions (p = 0.0020 BC/Aitchison, **Figure 3D**). Only the two extraction buffers led to comparable sample compositions (p = 0.35 BC/0.26 Aitchison, **Figure 3E**). These results in the even mock were independent of the chosen Bray-Curtis or Aitchison distance measure.

Analysis of the staggered mock (considering eight mock taxa, **Supp. Figure 6A-E**) and spike-in samples (considering three spike-in taxa, **Supp.Figure 6F-J**) overall confirmed the results of the even mock. We found significant differences in sample composition between lysis conditions in the staggered mock (p = 0.008 BC/0.21 Aitchison), and in the spike-in samples between all protocols (p = 0.012 BC/0.028 Aitchison) and between extraction kits (p = 0.012 BC/0.058 Aitchison). These larger and less conclusive p-values might be explained by the smaller sample size of only two instead of four dilutions per sample group.

Based on aforementioned results, we removed buffer contaminants and cross-contaminants of mock origin from the skin microbiome samples. Interestingly, the skin samples differed significantly only between the two subjects (p = 0.01 BC/0.18 Aitchison, **Supp.Figure 6K**). Apparently, differences in individual skin genera were so pronounced that they masked any other effect of extraction protocols in the skin samples (p > 0.074, **Supp. Figure 6L-O**).

### Which extraction protocol is best?

With substantial differences observed in sample composition between protocols, we aimed for determining the best extraction protocol. Therefore, we compared the sample composition of each cell sample to their corresponding DNA mock samples, in order to only evaluate extraction bias and to not bias our analyses, e.g., by differential amplification bias between taxa.

With lower distances and lower ranks indicating less extraction bias (**Supp.Figure 7A, B**), none of the protocols achieved a perfect representation of the expected DNA mock-based sample composition. Interestingly, although the choice of buffer did not significantly affect the sample composition, we found that each extraction kit consistently produced less bias when combined with its corresponding extraction buffer (**Supp. Figure 7C**). Again, input cell dilutions and the choice of BC or Ait distance measure led to very similar results.

In contrast, the performance of extraction protocols was very distinct between the even and the staggered mock community. Although good results were achieved for example with the Q_T_q and the Z_T_z protocol in both mock communities, it is impossible to determine a single best extraction protocol. But even more importantly, it seemed that the bias per protocol depends on the mock sample composition, in turn indicating that extraction bias does not only vary between protocols, but also between taxa.

### Morphology-based correction of extraction bias

To follow up the hypothesis of taxon-specific, protocol-dependent extraction bias, we used the compositional metacal approach [1] to calculate extraction bias per species and protocol. Bias in cell mocks was summarized across dilutions and compared to corresponding DNA mocks for even and staggered mock and to the expected composition for the spike-in community.

As hypothesized, we indeed found that extraction bias profiles varied substantially between species (**Figure 4**). However, some groups of species also presented very similar bias profiles over the eight extraction protocols, and even over different mock or spike-in sample compositions. We found that these groups were distinctively defined by the bacteria’s cell shape and cell wall structure (**Table 1**). Due to the small set of only eight species, we were not able to distinguish if the effect of cell shape is actually an effect of cell length/size, and continue using the terms ‘shape’ and ‘gram stain’ for simplicity. Still, bacterial cell morphology clearly seemed to determine protocol-specific extraction bias, independent of the sample composition.

Next, we tested if these protocol- and morphology-based bias profiles can be used for correcting extraction bias. Therefore, we calculated the extraction biases for eight species and eight protocols, but using only a single sample (10^6^ cell even mock, corresponding to “training data”, based on high sequence accuracy determined in previous sections). We summarized these 64 bias values by our morphology groups, leading to 24 correction factors (8 protocols * 3 morphology groups). These factors were then applied to correct new samples (corresponding to “validation data”). Success of bias correction was evaluated as a decrease in the distance to the DNA or expected mock composition.

A significant reduction of bias was achieved when correcting the 10^6^ even sample (p = 0.0078, internal correction of the training sample), but also when correcting validation dilutions of the even mock not involved in the bias calculation (p = 0.0078 for 10^8^ cells, p = 0.055 for 10^5^ cells, p = 0.0078 for 10^4^ cells, **Figure 5A**, **Supp. Figure 8A**). Correcting a different sample composition in the staggered mock did not significantly improve the bias (p > 0.31, **Figure 5B**, **Supp. Figure 8B**), but still reduced the median distance to the DNA sample composition. As a last step, we applied the bias correction to the spike-in samples, which represent different samples, with different sample composition, and even with completely different species as compared to the training sample. Here, the median distance to the expected composition was again reduced, reaching even a significant improvement in the 10^5^-cell sample (p = 0.012, **Figure 5C**, **Supp. Figure 8C**).

## Discussion

Microbiome sequencing data are distorted by errors and protocol-dependent biases, confounding biological interpretations. Using dilution series of well-characterized mock communities, we investigated sequence errors, chimeras, contaminants, and cross-contaminants. With the resulting precise species’ relative abundances, we revealed novel associations between extraction bias and bacterial cell morphology, which can be used to correct extraction bias bioinformatically.

Samples of the even mock community were characterized by particularly high proportions of sequence errors, i.e., ASVs with close genetic distance to expected sequences. The consistent and highly abundant presence of a few major sequence error ASVs across replicates would suggest they are true minor 16S gene variants not covered by the reference genomes. However, consistent detection does not prove sequence validity [23]. Instead, these ASVs may represent deviations from actual sequences that are particularly susceptible to error formation [39] and are not corrected by our denoising approach using DADA2 [40]. Independent of being minor variants or artifacts, most sequence error reads originated from the three expected species *E. coli*, *L. fermentum*, or *S. enterica*. Without accepting those sequence errors as true sequences, we would have critically underestimated the relative abundances of these three species in downstream analyses.

Interestingly, the presence of both sequence errors and chimeras increased with higher bacterial cell numbers. Assuming that higher cell numbers lead to higher DNA concentration and thus to closer physical proximity between similar DNA motifs from different species, high bacterial input would promote chimera formation by template switch during PCR amplification [41]. In line with that, chimera formation occurred mainly between closely related species (even mock) and, to a smaller extent, between highly abundant species (staggered mock), as observed in our data and previous research [23]. It has been established that high DNA input material can inhibit the PCR and lead to unspecific amplification, and that high numbers of PCR cycles should be avoided to reduce chimera formation [2, 41]. However, with substantial chimera formation starting at 10^8^ bacterial input cells in our data, chimeras may particularly hamper taxonomic annotation and diversity estimation in microbiome samples from high-biomass environments, such as stool [42].

The threat of contamination to low-biomass samples is well-known [15, 17, 18], yet internal cross contamination between samples is rarely investigated [16]. Our mock samples presented several skin associated genera like *Cutibacterium*, *Pseudomonas*, *Staphylococcus*, and *Corynebacterium* [25]. These four genera were found among the top 10 genera of our skin samples but have also been identified as common contaminants in microbiome sequencing experiments [17], e.g., originating from lab operators [15]. We also detected mock taxa in skin and control samples and found spike-in taxa (alien to the human microbiome and lab reagents [17]) in negative controls. Therefore, we conclude that our low-biomass samples are subject to considerable cross-contamination, which would have been inseparable from external contamination in environmental microbiome studies.

A major source of external contamination are bacterial DNA extraction kits, which have been previously identified to harbor a specific ‘kitome’ [15, 43]. Interestingly, most external contaminants in our data were not associated with extraction kits but with buffers. A previous investigation of contaminants in extraction kit components did not report buffer-specific contamination profiles but did not explicitly investigate external, non-mock taxa [44]. Validation of our findings would provide a simple roadmap to notably reduce contamination by treating only extraction buffers, e.g., with UV irradiation [45].

The two extraction buffers in our study lead to distinct contamination profiles, but apart from this, the buffers did not significantly alter the global sample composition of expected mock taxa, despite giving slightly less biased sample compositions when combined with their corresponding extraction kits. In contrast, the choice of extraction kit and lysis condition (speed and duration) significantly affected the relative abundances of mock taxa in our data. Although these differences were smaller than inter individual differences in the skin microbiome, observed here as well as in previous research [4, 46], extraction protocols still significantly contribute to the bias observed in microbiome sequencing data. Numerous studies have addressed this problem by benchmarking DNA extraction methods [47], usually aiming at determining the best extraction protocol for specific microbiome environments. In support of this research direction, we demonstrated that choosing the best protocol with the least extraction bias depends on the sample composition and taxa of interest.

However, no extraction protocol achieved perfect accuracy, each leading to more or less distorted sample compositions. Besides improving laboratory extraction methods, extraction bias could be resolved by a bioinformatic correction using defined bacterial mock communities. Here, we provide novel evidence for such a bioinformatic correction by linking extraction bias per protocol and taxon to bacterial cell morphology, specifically gram stain and cell shape. It is well-established that gram positive bacteria are harder to lyse than gram-negative bacteria, demanding a mechanical lysis step such as bead-beating to lyse the thick gram-positive cell wall. In the context of cell lysis, gram stain only serves as a proxy for cell wall thickness. For example, *Truepera radiovictrix* cannot be classified into gram-positive or gram-negative stain but possesses a thick three-layered cell wall [48] that led to similar extraction efficiencies like other gram-positive bacteria. Similarly, the effect of cell shape on extraction bias could have been equally represented by bacterial cell size. In lack of support for either hypothesis, we propose that rod-shaped (= larger) bacteria are more easily hit by beads during mechanical lysis. In line with that, gram-negative and rod-shaped bacteria seemed to be rather overrepresented compared to gram-positive or coccoid bacteria, and the tougher protocol (‘T’) slightly enhanced the relative abundance of gram-positive cocci.

Using the morphology-based bioinformatic correction, we significantly reduced extraction bias in new samples and even in new taxa using gram stain and cell shape. Additional factors, such as aerobic status, motility, or spore formation, may affect cell stability in the lab and could have improved our bias correction model. However, we were limited to a sparse set of factors in our 8-species mock community. Interestingly, previous research has claimed that bias cannot be summarized for groups of taxa [1], supported by differential biases observed between species of the same genus [1, 2, 49]. However, the mock experiment by Brooks et al. [50] used species without available reference genomes, leading to ambiguous estimations of 16S copy numbers [1, 50] and potential problems with taxonomic annotation, which jointly blur precise abundance estimations. The experimental design by Morgan et al. [49] did not allow for specifically measuring extraction bias. Thus, differential amplification may have affected the bias of closely related species [1, 50]. We believe that only our combination of well-characterized mock communities with reference genomes, rigorous study design including corresponding DNA mocks, and in-depth analyses of downstream biases allowed for revealing an association between bacterial cell morphology and extraction bias.

Extraction bias will remain a major confounder for 16S sequencing experiments and will not be alleviated by recent advances in sequencing technologies, such as long-read sequencing or shotgun metagenomics. To date, extraction bias is only addressed by consistently using the same lab protocol to keep the bias equally constant [2]. Instead, we found a novel association between extraction bias and cell morphology, which may enable the bioinformatic correction of extraction bias using standardized controls. Our findings pave the road for cross-protocol meta-analyses and for discovering more robust clinical microbiome associations.

## Acknowledgements

We thank our study participants and ZIEL for sequencing. The ZymoBiomics extraction kit was kindly provided by ZymoResearch.

---

**Supp. Figure 1.**
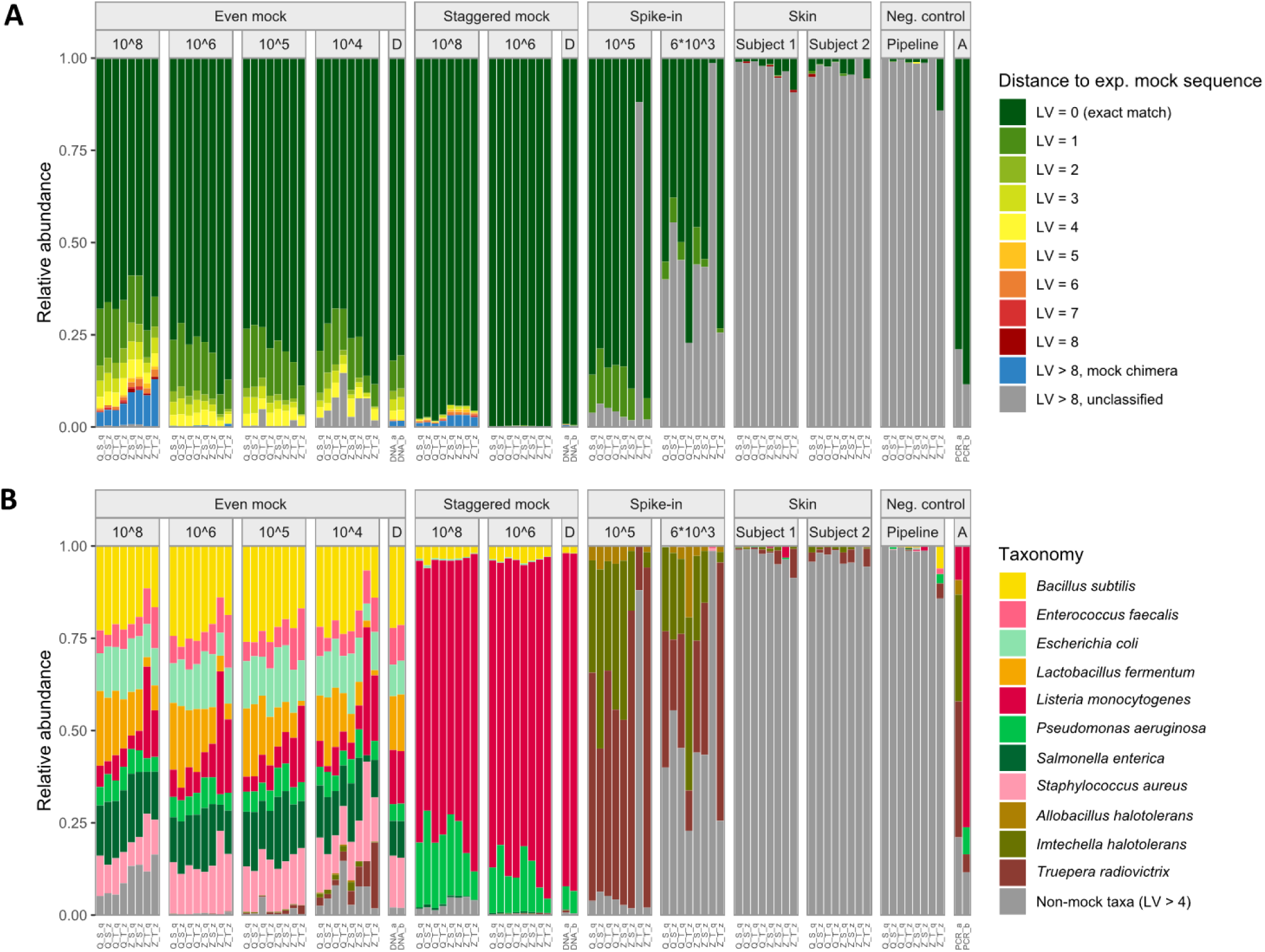
Taxonomic annotation by comparison to expected sequences reveals sequence errors, chimeras, contamination, and differences in sample composition between extraction protocols. Levenshtein (LV) distance between observed ASV sequences and expected mock sequences allows for classifying sequences into exact matches, sequence errors, chimeras, and remaining ‘unclassified’ sequences with LV > 8 (**A**). After accepting sequence errors with LV ≤ 4 as correct ASVs, considerable differences in sample compositions between extraction protocols were found, but also cross-contamination from mock into skin and negative controls samples (**B**). Chimeras (**A**) were defined as sequences with LV > 8 to any expected sequences, and > 95% identity by LCS with at least two expected mock sequences. Sample composition (**B**) is shown for the 11 expected mock taxa, with non mock taxa (LV > 4) representing, e.g., contaminants and skin taxa. Q: Qiagen extraction kit, Z: ZymoResearch extraction kit, S: ‘soft’ lysis condition, T: ‘tough’ lysis condition, q: Qiagen/Stratec buffer, z: ZymoResearch buffer. A: Amplification/PCR control. D: DNA mock samples.

**Supp. Figure 2.**
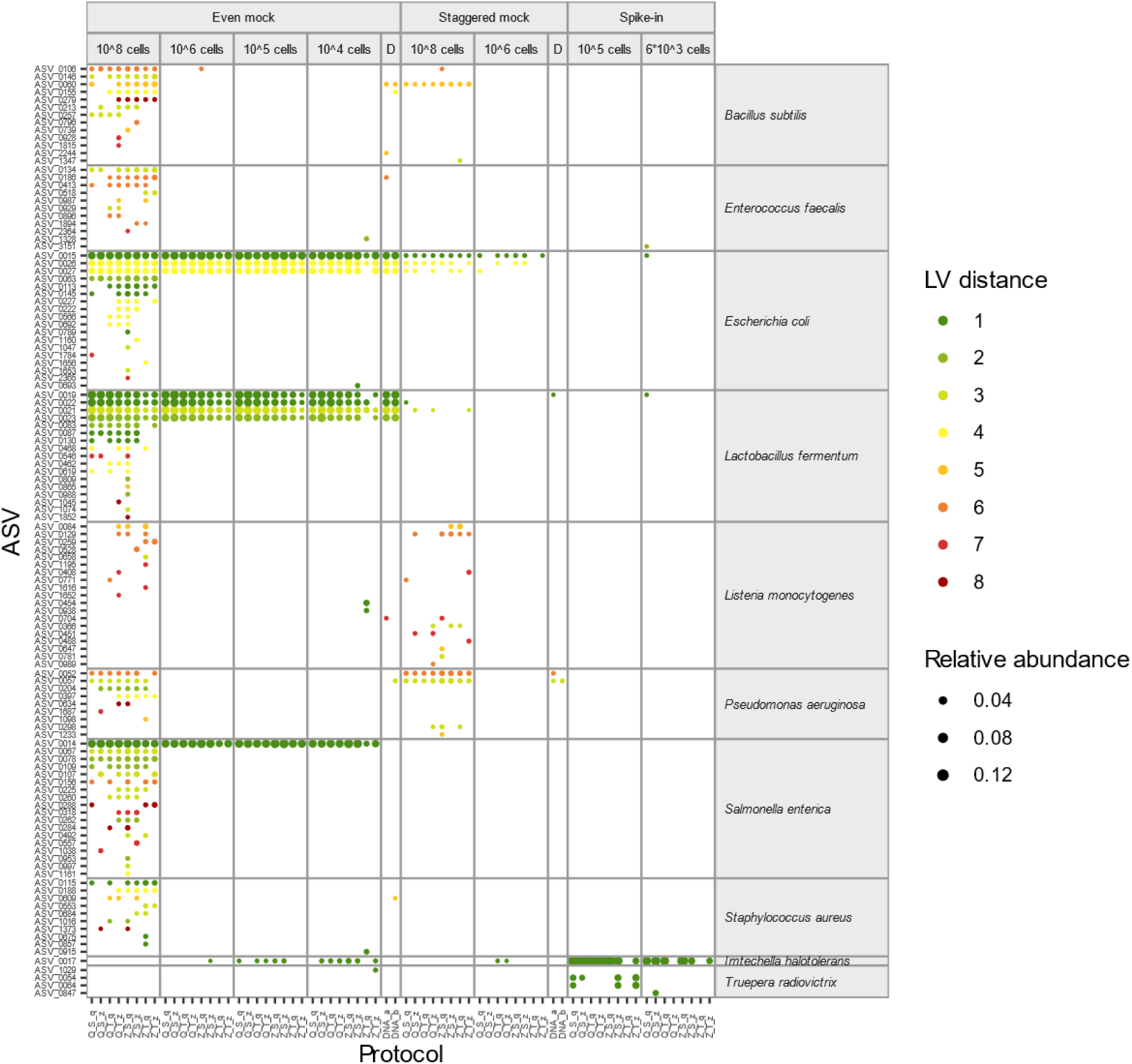
Sequence errors are mostly assigned to *E. coli*, *L. fermentum*, or *S. enterica*, independent of the extraction protocol. Sequence errors were classified as ASVs with Levenshtein (LV) distance ≥1 and ≤ 8 to expected mock sequences. Point area indicates ASV relative abundance per sample. D: DNA mock sample.

**Supp. Figure 3:**
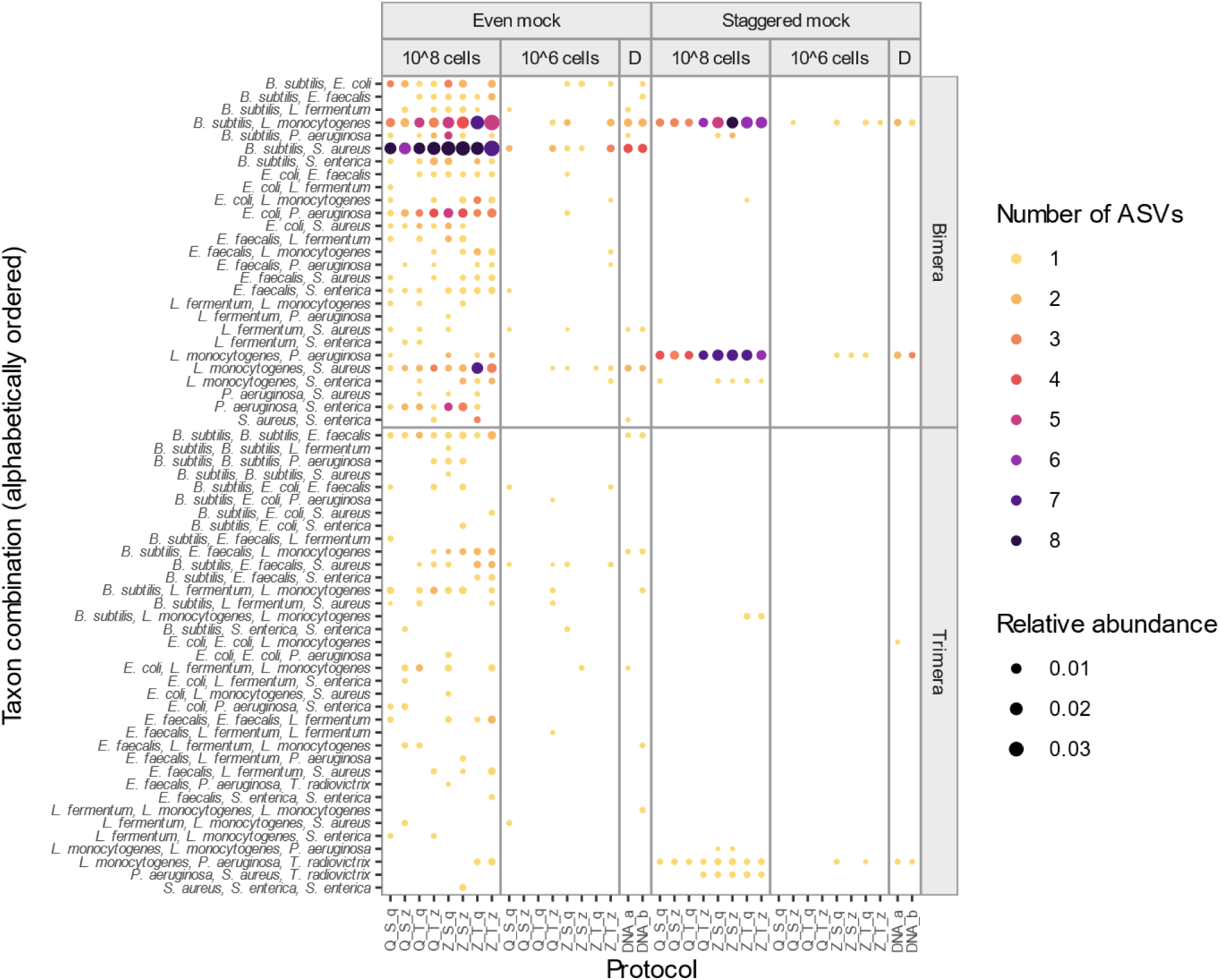
Chimeras are predominantly formed in high-input cell samples, independent of the extraction protocol. Bimera and trimera taxon combinations are only shown for samples with 10^8^ or 10^6^ input cells of the even and staggered mock community. Chimeras were defined as ASVs with Levenshtein (LV) distance ≥ 8, and > 95 % sequence identity with at least two expected mock taxa. Point area indicates each chimera combination’s relative abundance per sample. D: DNA mock sample.

**Supp. Figure 4.**
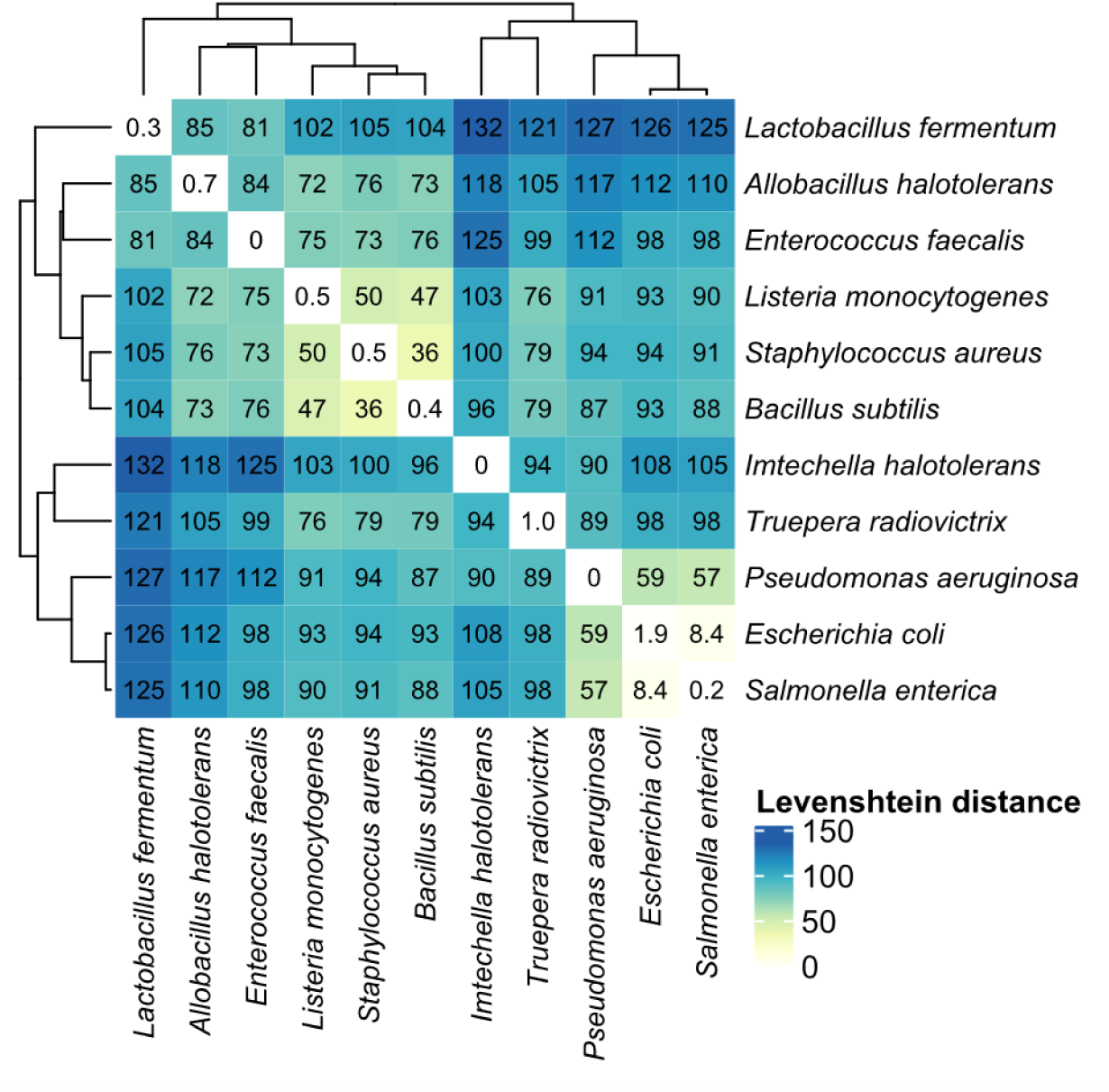
Sequence distances between mock and spike-in expected sequences. Levenshtein (LV) distance specifies the number of substitutions or indels between sequences, with zero indicating identical sequences. Clustering analysis of mean LV distances highlights closely related expected sequences between *E. coli*, *S. enterica*, and *P. aeruginosa*, and between *S. aureus*, *B. subtilis*, and *L. monocytogenes*. Values represent mean LV distances between 16S rRNA copy variants of two species, based on reference sequences provided by ZymoResearch and cut to 279 bp of the V1-V3 region. Values in the diagonal indicate mean LV distances between copy variants within each species. Clustering was performed with Euclidean distance and complete linkage.

**Supp. Figure 5.**
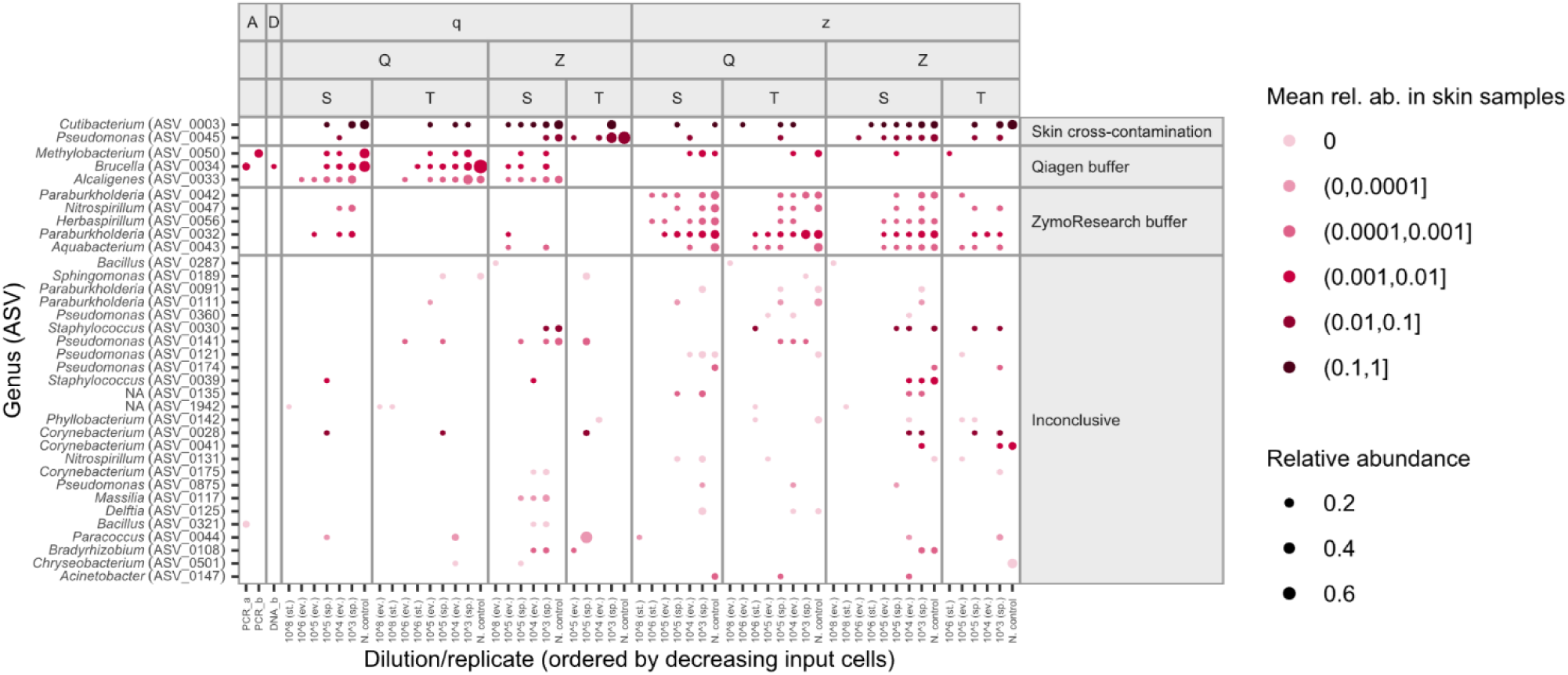
Most contaminant reads in mock samples and negative controls originate from cross contamination of skin samples or from extraction buffers. Contaminating ASVs were clustered into four groups by kmeans. Clusters were assigned to buffer origin by their distinct and consistent appearance across samples of the same extraction buffer, and to skin origin by their high relative abundance in skin microbiome samples (indicated by darker color). Shown are ASVs previously categorized as ‘Unclassified’, i.e. with LV distance ≥ 8 and ≤ 95 % identity with expected mock taxa sequences, and present in at least three non-skin samples (mock or negative controls). Point area indicates ASV relative abundance per sample.

**Supp. Figure 6.**
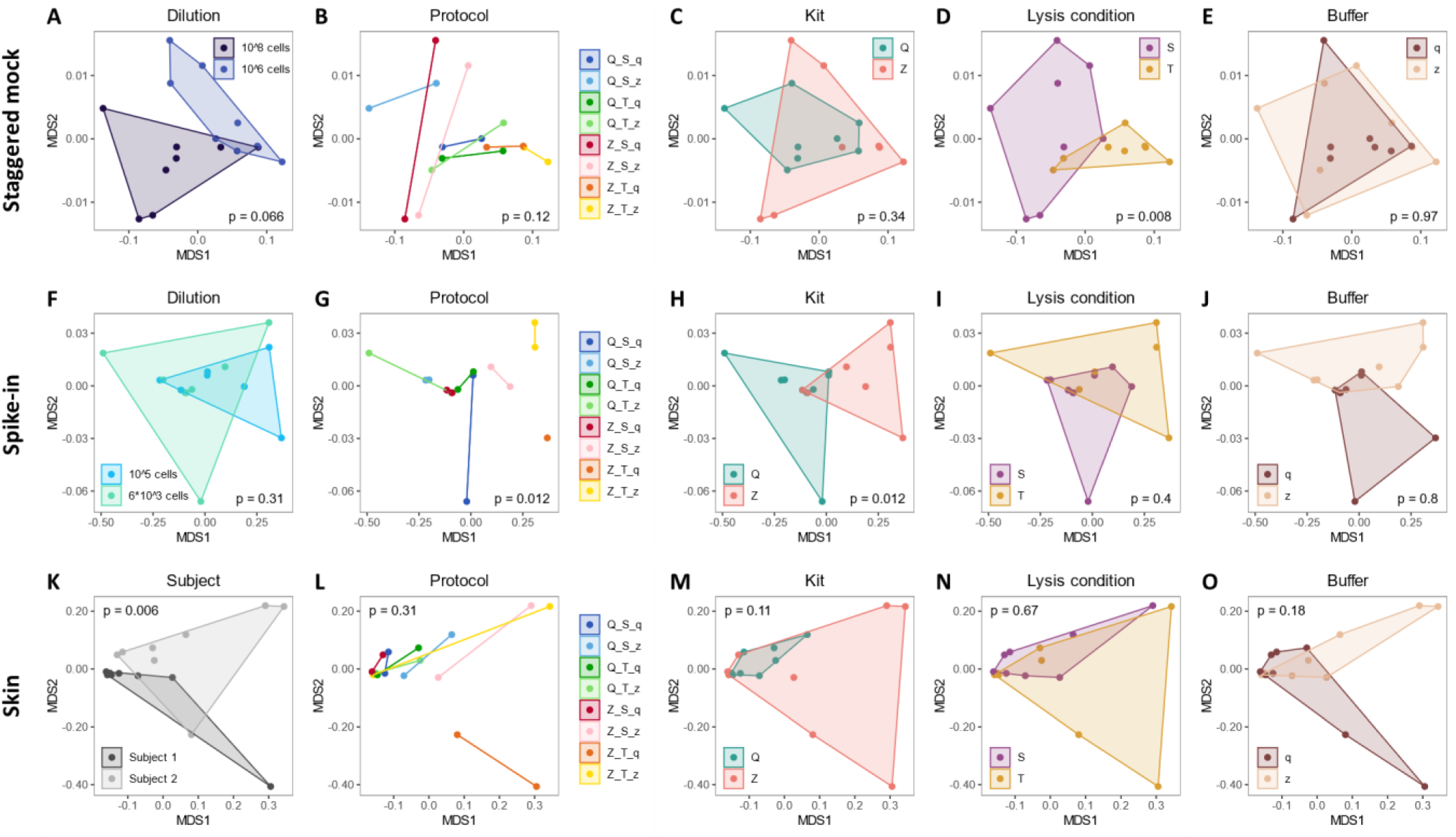
Sample composition in the staggered or spike-in mock community is significantly affected by extraction protocols, kit, or lysis condition, but microbiome compositions between two subjects are even more distinct than between extraction protocols. In the staggered mock community (**A-E**), beta diversity analysis revealed significant differences in global mock composition between lysis conditions (**D**), but not between dilutions (**A**), protocols (**B**), kits (**C**), or buffers (**E**). In the spike-in community (**F-J**), protocols (**G**) and kits (**H**) significantly affected sample compositions, but not dilutions (**F**), lysis conditions (**I**), or buffers (**J**). In contrast, in the skin microbiome samples (**K-O**), significant differences in global microbiome composition were only detected between the two subjects (**K**), but not between any of the extraction protocol variables (**L-O**). Beta diversity was performed only on mock taxa with LV ≤ 4 to any expected mock sequence, and is visualized by PCoA on Bray-Curtis dissimilarities. Polygonal shaded areas connect samples of the same group, p-values are derived from PERMANOVA tests with 500 permutations. Q: Qiagen extraction kit, Z: ZymoResearch extraction kit, S: ‘soft’ lysis condition, T: ‘tough’ lysis condition, q: Qiagen/Stratec buffer, z: ZymoResearch buffer.

**Supp. Figure 7.**
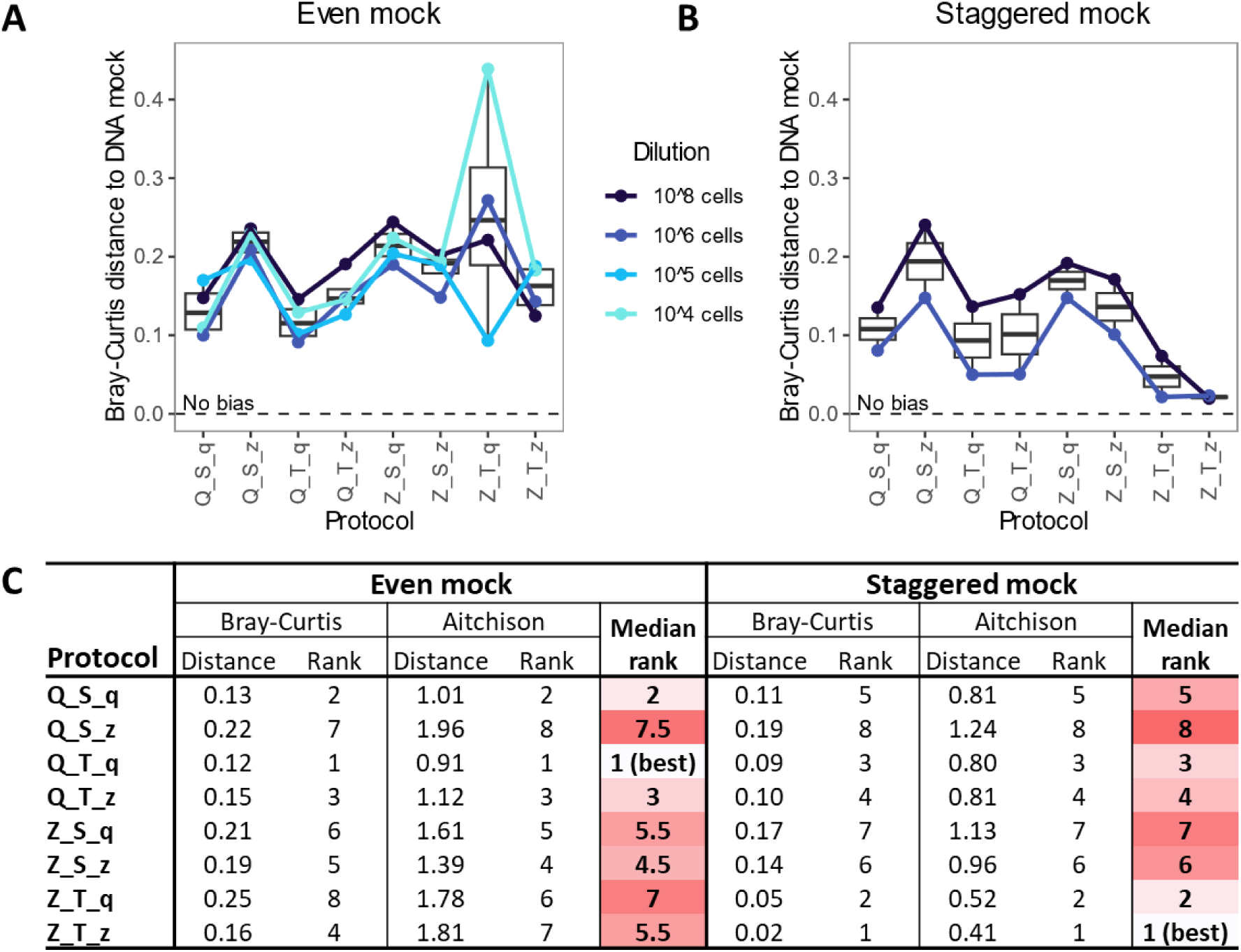
Magnitude of extraction bias varies between extraction protocols, but also between mock communities. Extraction bias, measured as Bray-Curtis distance to the DNA mock composition, varies between extraction protocols in the even (**A**) and staggered (**B**) mock community, with no protocol achieving a perfect representation (no bias). Substantial differences in protocol bias were observed between the two mock communities, independent of the chosen distance measure (**C**). Boxes (**A**, **B**) denote the median and interquartile range (IQR), whiskers represent values up to 1.5 times the IQR, dots indicate individual samples. Darker red background color (**C**) indicates higher extraction bias per protocol.

**Supp. Figure 8.**
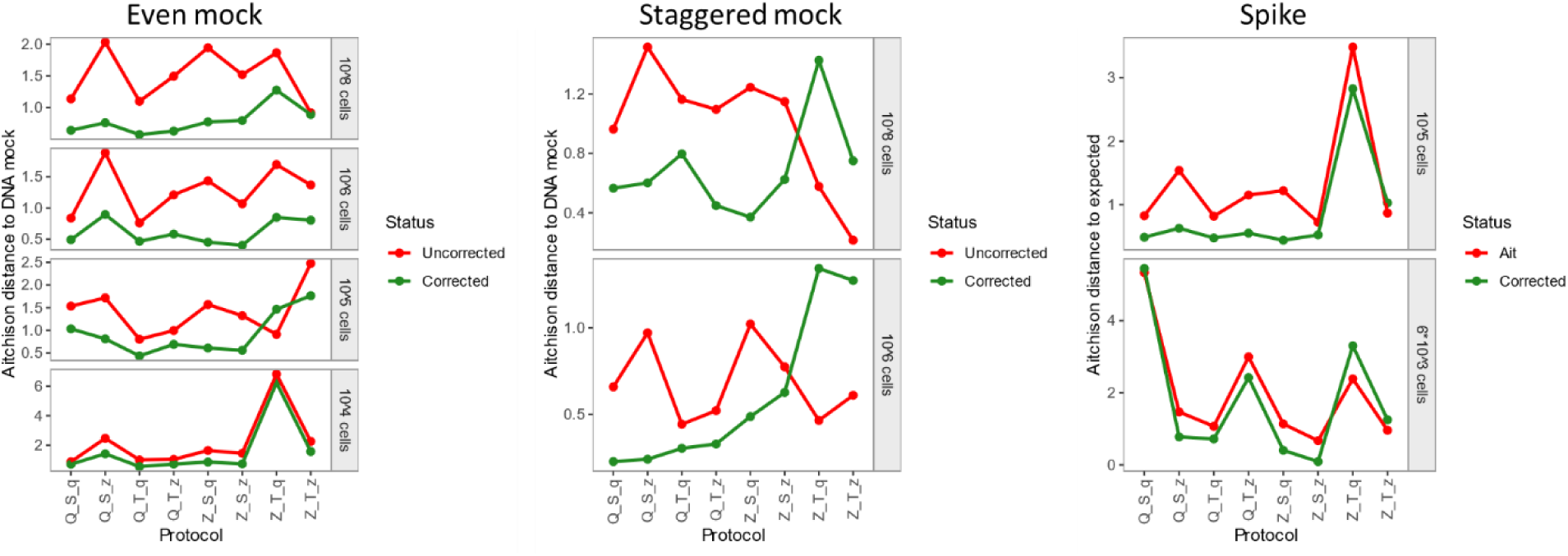
Distance to the DNA or expected mock composition is reduced in most protocols after applying morphology-based correction of extraction bias. Extraction bias per protocol was calculated from the 10^6^ even mock samples, summarized by bacterial morphology group, and applied to the 10^6^ even mock sample (internal correction), but also to different samples of the same mock (even mock 10^8^, 10^5^, 10^4^ cells), different samples of a different mock (staggered mock), and to different samples with different taxa (spike-in mock). Extraction bias was measured as Aitchison distance to the DNA mock composition (even mock, staggered mock) or as distance to the expected mock composition (spike-in mock).

